# Analysis of the three dimensional structure of the kidney glomerulus capillary network

**DOI:** 10.1101/677864

**Authors:** Mark Terasaki, Jason Cory Brunson, Justin Sardi

## Abstract

The capillary network of the kidney glomerulus filters small molecules from the blood. The glomerular 3D structure should help to understand its function, but it is poorly characterized. We therefore devised a new approach in which an automated tape collecting microtome(ATUM) was used to collect 0.5 micron thick serial sections from fixed mouse kidneys. The sections were imaged by scanning electron microscopy at ∼50 nm / pixel resolution. With this approach, 12 glomeruli were reconstructed at an x-y-z resolution ∼10x higher than that of paraffin sections. We found a no-cross zone between afferent and efferent branches on the vascular pole side; connections here could allow blood to exit without being adequately filtered. Network analysis indicates that the glomerular network does not form by repetitive longitudinal splitting of capillaries. It also suggests that capillaries vary their diameter to make flow more efficient. The shortest path (minimum number of branches to travel from afferent to efferent arterioles) is relatively independent of glomerular size and is present primarily on the vascular pole size. This suggests that the shortest path is established on the vascular pole side, after which new branches and longer paths form on the urinary pole side. Thus the 3D structure of the glomerular capillary network provides useful information with which to understand glomerular function. Other tissue structures in the body may benefit from this new three dimensional approach.

## Introduction

There are approximately 1 million nephrons in a human kidney (e.g. Eaton and Pooler, 2018). Their collective function is to maintain the homeostatic concentration of ions and small molecules in the blood. The first part of the nephron is the glomerulus, a capillary “tuft” enclosed within Bowman’s capsule. The capillary network performs the initial filtration of water, ions and small molecules from the blood. In a healthy person, 20% of the cardiac output goes to the kidneys, of which 10-15% is filtered by the glomerulus (the glomerular filtration rate). The filtrate passes through the tubular system of the nephron, where needed materials are resorbed and ion concentrations adjusted, leaving toxic or uneeded substances which will be excreted. Glomeruli are directly damaged by auto-immune diseases or toxic chemicals and are secondarily damaged in diseases such as diabetes and hypertension. Severe damage leads to end-stage renal disease (ESRD), a leading cause of death.

A single input (afferent) arteriole feeds into the glomerulus and a single output (efferent) arteriole leads away from it. The afferent arteriole branches to form a complex network of capillaries that converges back to the efferent arteriole. It has been difficult to document the exact branching pattern of the capillaries. Paraffin sections, the standard histological method, are typically 4-5 microns thick or thicker. This is not thin enough to reconstruct the capillaries in three dimensions. The glomerular network has been imaged by x ray nanotomography (Wagner et al., 2011), but the exact branching pattern in the interior was not determined. The glomerulus has been reconstructed by embedding in epon, collecting 1 micron thick serial sections and imaging by light microscopy with a high numerical aperture objective lens (Remuzzi et al., 1992). This was so challenging that only one glomerulus was analyzed. Other studies have similarly reconstructed only one or have done partial reconstructions (see Discussion). Thus, the capillary network has not yet been adequately characterized in three dimensions.

Recently developed techniques for serial section electron microscopy offer new ways to investigate the three dimensional structure of biological structures. This is made possible by automated collection, as well as radical improvements in computer speed and software for alignment and 3 dimensional analysis. We have adapted the ATUM method (Kasthuri et al., 2015) for the glomerulus, and have reconstructed 12 glomeruli. Network analysis (classically known as graph theory) can be very useful for analyzing glomerular connectivity (Wahl et al., 1984; Wahl et al., 2004), and we used some of these tools to analyze glomerular properties.

## Results

### 500 nm sections

The ATUM tape collector method picks up sections on tape and the sections are imaged by a scanning electron microscope. Using this method, 2000 sections, each 30 nm thick, were cut and collected from mouse brain cortex (Kasthuri et al., 2015). A 50 *µ*m x 50 *µ*m field of view was imaged at 3 nm per pixel resolution (16,000 x 16,000 pixel images).

A similar approach could be used on the mouse glomerulus, which is approximately 70 microns in diameter. However, even a single glomerulus would be a major undertaking due to the difficulty of collecting several thousand sections and lengthy imaging time. We realized that cutting thicker sections and imaging at lower resolution would still provide enough information to reconstruct the glomerular capillary network, and would allow us to image multiple glomeruli to compare their structures.

For imaging by transmission electron microscopy, sections must be thin enough so that electrons can pass through, typically 60-70 nm. With scanning EM, the image signal is formed from electrons that are scattered back from the superficial layers of the section surface. Thus, the surface of sections of arbitrary thickness is easily imaged. A 70 *µ*m diameter glomerulus cut into 0.5 *µ*m thick sections requires only 140 sections. Although high resolution imaging (e.g. 5 nm per pixel) is somewhat compromised with thick sections, a moderate image resolution of 50-100 nm per pixel (which is considered to be “super-resolution” by light microscope standards) is adequate to resolve cell boundaries and many tissue features.

Serial section images of the mouse glomerulus show the capillary network clearly (Figure 1). The arterioles at the vascular pole as well as the beginning of the proximal tubule at the urinary pole are easily identified. The afferent arteriole entering the glomerulus can be identified because it branches from a large artery, whereas the efferent arteriole dives down into the medulla of the cortex. We collected serial sections of 12 glomeruli from 4 different mice. For each, the capillary lumen was segmented, reconstructed and rendered (Figure 2).

**Figure 1.**
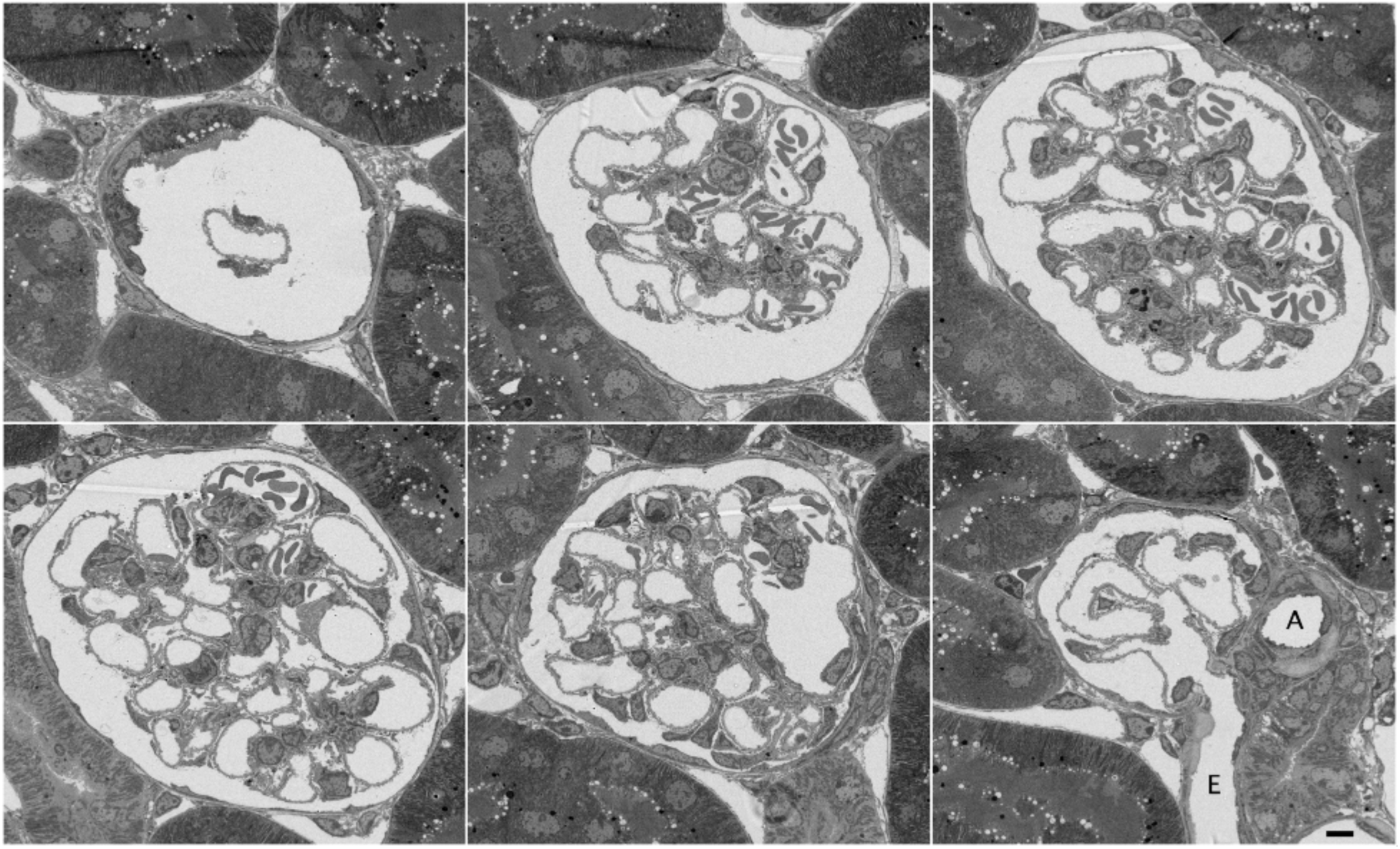
Montage of serial section electron micrographs. The sections were 500 nm thick, and were imaged at 66 nm per pixel. The section numbers were 30, 55, 80, 106, 137 and 159 out of 238 total sections. “A” marks the afferent arteriole, and “E” denotes the efferent arteriole. These came from glomerulus #6 of Figure 2. Scale bar = 5 *µ*m.

**Figure 2.**
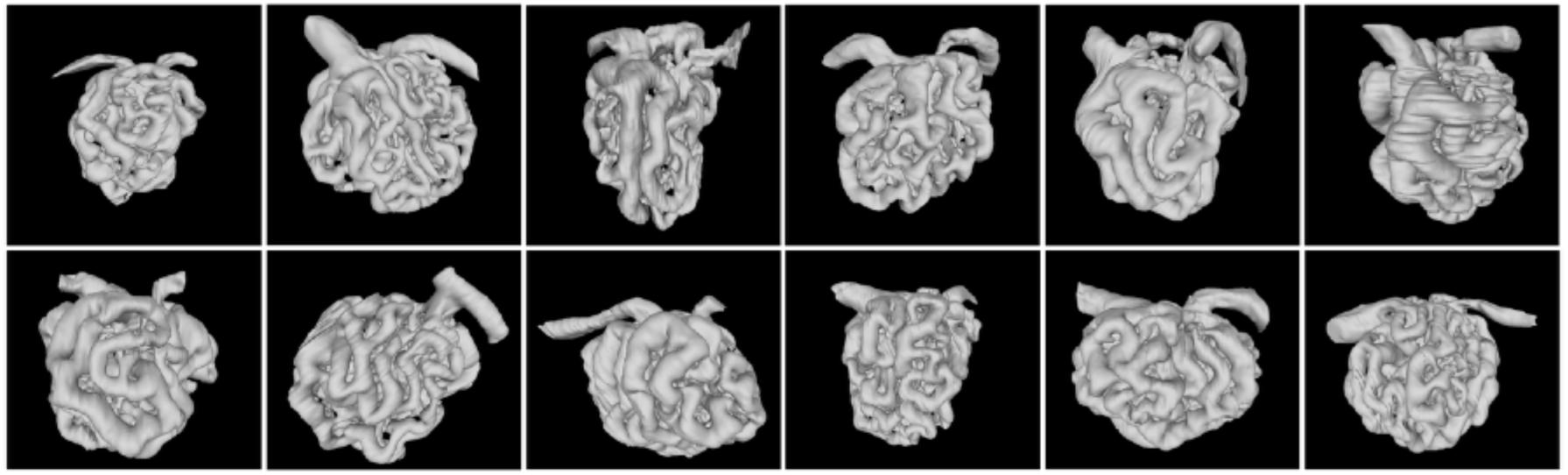
Montage of reconstructions of 12 glomeruli. Serial section data was segmented then 3D models were generated. They are shown from smallest (fewest nodes) to largest. The afferent and efferent arterioles are oriented up with the afferent arteriole on the left.

Certain features are evident on visual inspection of the reconstructions. It is commonly stated that the afferent branches are wider than the efferent branches, corresponding to the net 10-15% loss of volume as the blood travels through the glomerulus. The reconstructions show this clearly (Figure 3).

**Figure 3.**
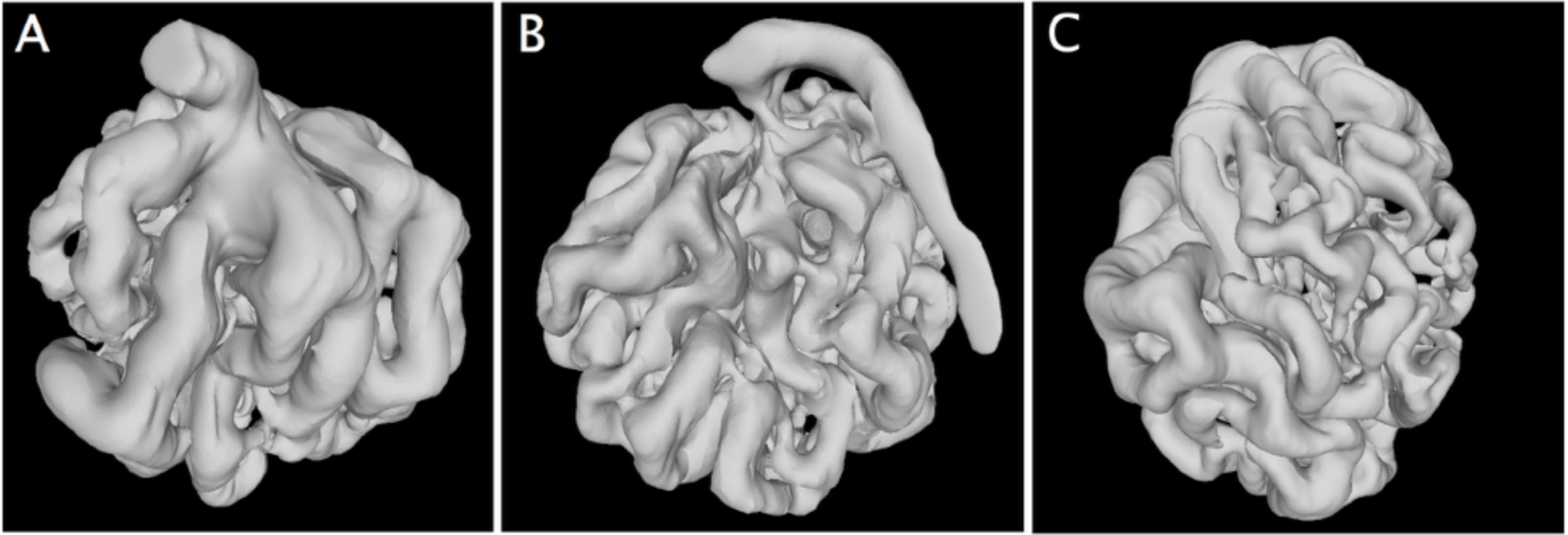
Views from opposite sides of a glomerulus reconstruction showing that the afferent arteriole branches (A) are noticeably wider than the efferent arteriole branches (B). This is from the glomerulus shown in the bottom row, far left of Figure 2. C) View from the urinary pole of the same glomerulus.

### Lack of short cuts from afferent to efferent branches

When the reconstructed glomeruli are viewed from the vascular pole side (i.e., the side with the two arterioles), there is invariably a gap which separates the afferent and efferent arteriole branches (Figure 4). This is consistent with the function of the glomerulus because a connection in this region would provide a path for blood to exit without being filtered.

**Figure 4.**
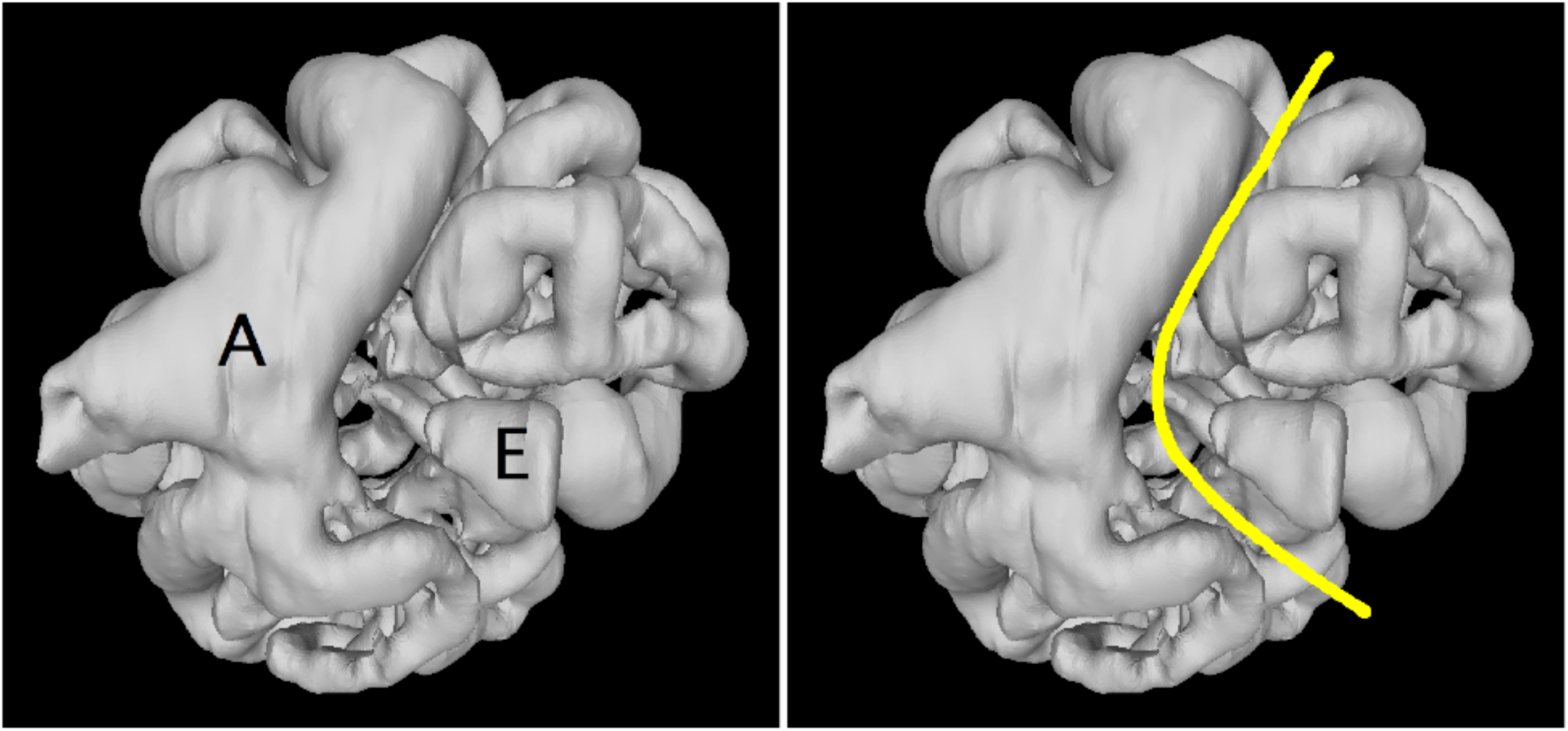
View of a glomerulus reconstruction from the vascular pole side, showing the branches from the afferent arteriole (“A”) and efferent arteriole (“E”). There are no connections between these branches visible on this side. They would constitute a short cut path which would allow blood to pass through without being filtered. Right panel shows the exclusion zone. As in this example, the afferent branch is larger usually.

During development, how are such connections prevented? It occurred to us this that this could be a consequence of longitudinal splitting of capillaries to form branches (e.g. Karthik et al., 2018). The capillary nework is thought to begin as a single loop which then branches (Quaggin and Kreidberg, 2008). There are 3 ways to add branches (Risau and Flamme, 1995; Risau, 1997): coalescence of mesenchyme cells (vasculogenesis), sprouting (angiogenesis), and splitting (intussusception) (Figure 5). Additions to a loop by repeated splitting forms a network with no short cuts. Repeated splitting also produces a “flat” network, i.e., one that can be drawn on paper without any crossing lines. As described below, this property can be used to determine whether a capillary network was generated by repeated splitting.

**Figure 5.**
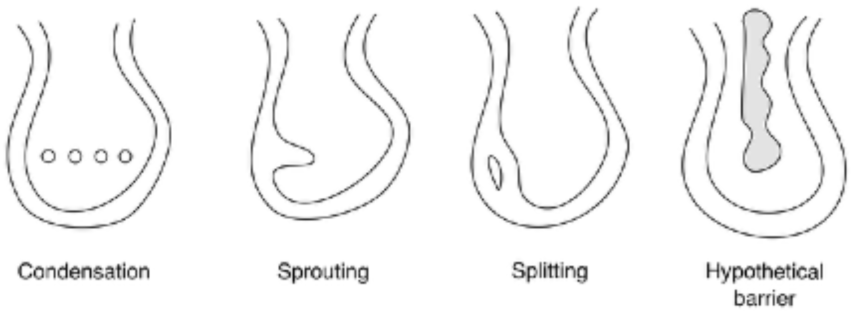
Three different ways to form capillary branches. From left to right: Condensation of endothelial cell precursors, also called vasculogenesis; Sprouting from pre-existing capillary; Widening of a capillary region following by longitudinal splitting. Far right: Hypothetical barrier that could prevent formation of capillary branches which would short circuit the glomerulus.

For network analysis, it is necessary to identify the nodes (intersection points) and how they are connected by edges (capillary branches). It was unexpectedly difficult to do this from the original serial section images. Essentially, the network is viewed from only one angle, and this makes it easy to mis-identify branch points or lose track of the path of the branches. We used the reconstructions (.obj files) instead of the image data, and viewed them with virtual reality optics which allowed us to rotate and slice the glomerulus at different angles. This enabled us to place nodes and connections within the reconstruction much more reliably and efficiently (Figure 6). When it was completed, the software exported a list of nodes, their positions, and their interconnections (“adjacency list”) necessary for network theory analysis.

**Figure 6.**
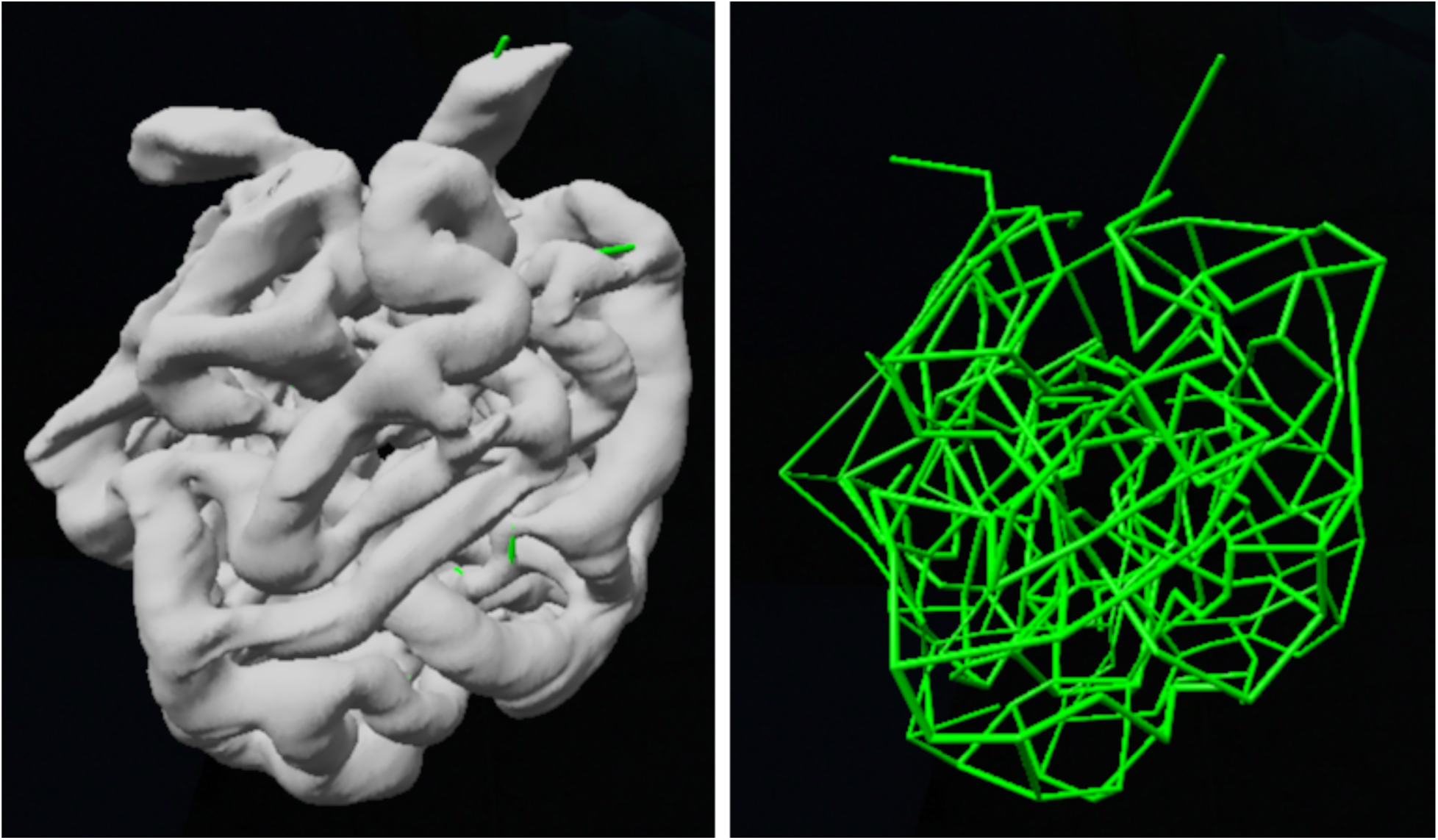
Three dimensional network diagram constructed from a glomerular reconstruction. The nodes and connections were identified and constructed in a 3D virtual reality system.

An algorithm that detects planar graphs using the Kuratowski theorem (e.g., Trudeau, 1994) shows that all 12 glomeruli are not flat. A similar conclusion was reached by an earlier study which analyzed 6 glomeruli from several different publications (Wahl et al., 2004). This is evidence that repeated longitudinal capillary splitting is not the sole mechanism for preventing short cuts nor for constructing the glomerulus network, though it is certainly possible that splitting may occur in parallel with sprouting or condensation. We propose another mechanism for preventing short cuts that involves a physical barrier inside the initial glomerular loop (see Discussion).

### A network analysis of flow

A standard network technique uses Kirchoff’s electrical circuit laws to model flow through a network of resistors (Newman, 2010). Kirchoff’s laws state that the current sums to 0 at each node, and that the voltage difference in any loop is 0. Circuit analysis uses standard linear algebraic (matrix) methods on a system of N equations and N unknowns to solve for the voltages at nodes and currents along the connections. This approach can be used for the glomerulus, in which pressure is analogous to voltage and flow is analogous to electric current. We set the resistance of each branch to be equal and used boundary conditions of fixed current in and out of the afferent and efferent arterioes. This is unrealistic, since the resistance to flow is proportional to capillary diameter, length and other factors. However, it is still potentially useful because the solution represents the flow which is determined solely by the network topology.

The results of one of these calculations is shown in Figure 7. The flow is lowest for branches on the urinary pole side, mid-way between the afferent and efferent arterioles. The diameter of the capillaries in this region are always smaller (e.g. Figure 3C). The idealized model which takes only topology into account predicts flows that are similar to the observed capillary diameters. This is consistent with the idea that the capillaries adapt to flow pattern that is determined by the topology (see Discussion).

**Figure 7.**
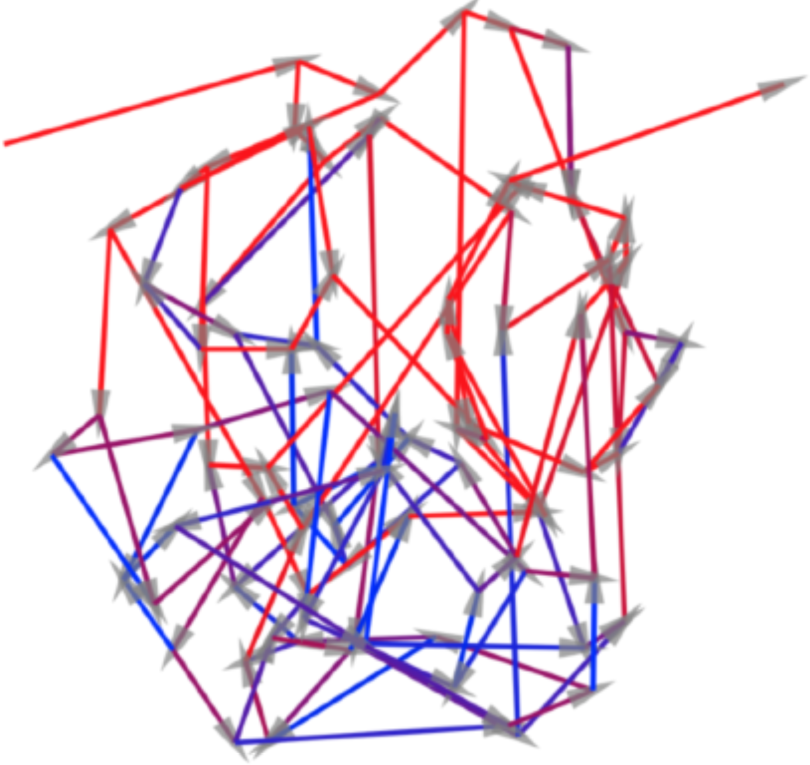
Flow direction and magnitude in a simplified model of a glomerulus. The glomerulus shown on the top row, far right of Figure 2 was analyzed using Kirchoff’s circuit laws with a voltage / pressure of 1.0 at the afferent arteriole and −1.0 at the efferent arteriole. The resistances of all branches was set to 1.0, so the calculation reports the current / flow where only the topology of the network is taken into account. Direction is indicated by arrows, and magnitude is indicated by color (red is high flow, blue is low flow). The flow magnitude decreases as the branches diverge from the afferent arteriole and then increases as the branches converge on the efferent arteriole.

### Differences between small and large glomeruli

An important network parameter is the shortest path, that is, the minimum number of nodes traversed by a path from the afferent arteriole entry to the efferent arteriole exit. We determined this using standard network methods (Newman, 2010). The number of steps in the shortest path ranged from 8 to 12. When plotted against the number of nodes, the shortest path is largely independent of the number of nodes (Figure 8A). Wahl et al (2004) made a similar observation from their analysis of 6 glomeruli from different studies.

**Figure 8.**
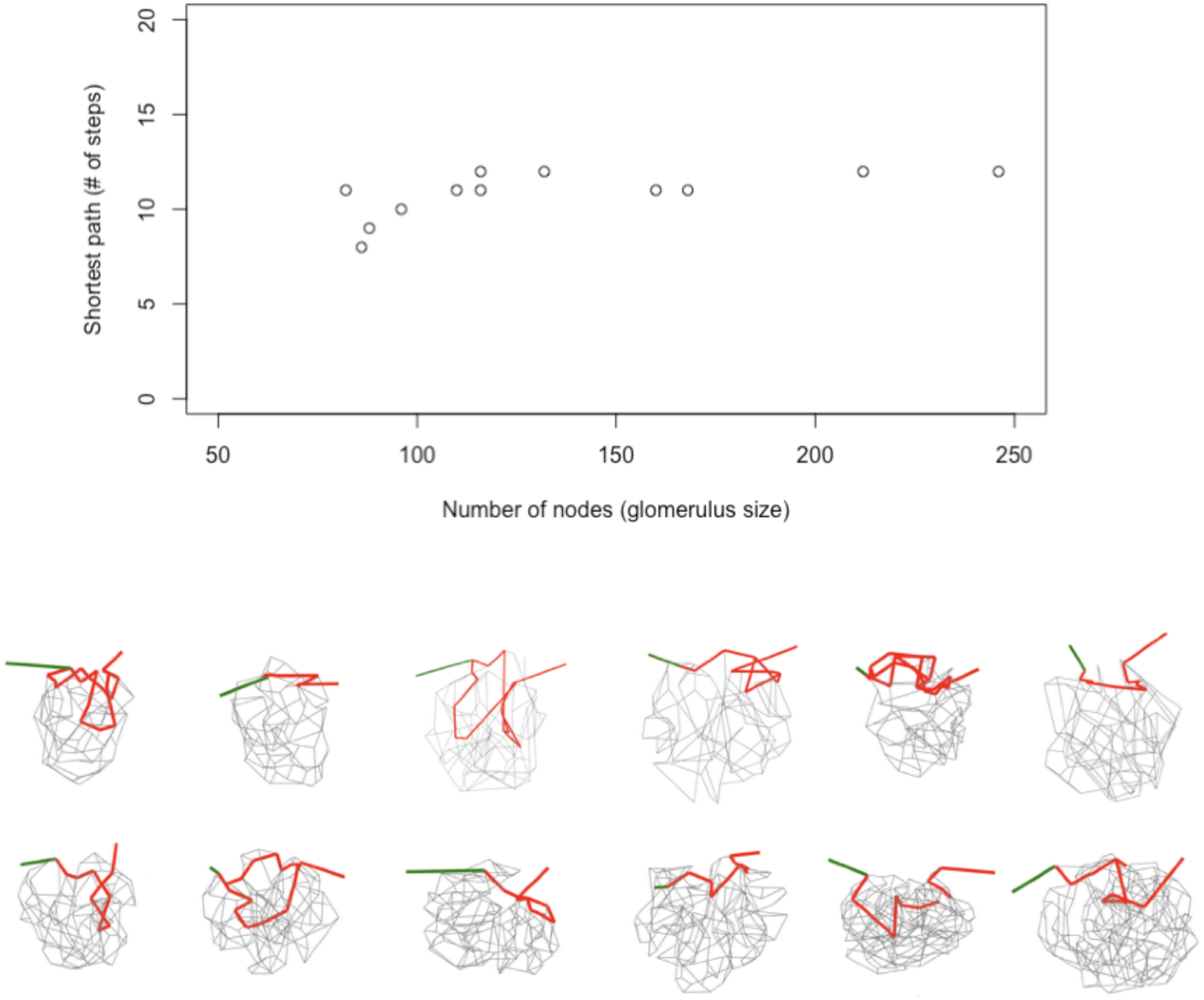
Shortest path versus glomerulus size. A) Graph of the number of steps in the shortest path versus the number of nodes (branch points) in the glomerulus. B) The glomeruli are shown in the same order as in Figure 2, and oriented similarly with the afferent arteriole on the left colored green. The shortest path(s) (in several instances, there are two paths of the same length) from afferent to efferent arteriole is shown in red, and the other connections within the glomerulus are shown in gray. The shortest path is located on the vascular pole side of the glomerulus. These observations suggest that the shortest path is established at some point during glomerular development, after which more branches are added on the vascular pole side which create paths of longer lengths.

When the shortest path(s) (sometimes there are two paths of the same length) is plotted, it is present in on the vascular pole side of all of the glomeruli (Figure 8B), avoiding the no-cross zone. This and the relatively constant number of steps in the shortest path suggests the shortest path is established at some point of glomerulus development, and that after this time, branches are added on the urinary pole, creating paths of longer lengths.

## Discussion

Why is the glomerulus a branched network, not a single long convoluted capillary? Two explanations, not necessarily exclusive, are redundancy and filtration efficiency. A branching network provides redundant pathways that could circumvent damage or an occlusion. A branching network also could filter more efficiently by exposing the blood to a greater amount of surface area and / or increasing the volume that can be fitered per unit time.

Since the glomerulus is in fact a branched network, does the exact type of branching pattern have significance? The flow biophysics can be very complex, added to the the fact that the pressure in the glomerular capillaries is higher than in the peripheral capillaries (Skorecki et al., 2012).

Questions like these could be better addressed if the connectivity of the glomerular capillaries were known. A number of studies have therefore been devoted to obtaining reconstructions (Shea, 1979; Shea and Raskova, 1984; Winkler et al., 1991; Remuzzi et al., 1992). Most recently, careful partial reconstructions of human glomeruli have been made (Neal et al., 2018). Advances in computer capabilities facilitate making reconstructions, and network theory is well suited for analyzing the connectivity (Wahl et al., 2004).

We have used a new approach involving serial section electron microscopy to fully reconstruct 12 mouse glomeruli. Visual inspection of the reconstructions verifies that the afferent branches are thicker than the efferent branches. There is a loss of fluid volume on passage through the glomerulus. This seems to be consistent with an adaptation of the capillary diameter to the amount of flow (discussed below).

### Lack of short cuts from afferent and efferent branches

Visual inspection of the reconstructions also shows that there is a no cross zone between the afferent and efferent branches on the vascular pole side of the glomerulus. This is an important feature of the glomerulus because connections in this region could provide a short cut for blood to pass through without filtration. As explained in the Results, this could be due to branch formation by repeated capillary splitting. Network analysis shows that the actual glomerular connectivity is not “planar”. This eliminates repeated splitting as the developmental process. Based on a more limited sample size, Wahl came to the same conclusion regarding planarity of glomerular networks, though they did not consider the implications for network formation.

Studies of the early development of the glomerulus suggest another way for preventing short cuts between afferent and efferent branches. The initial structure forms an S shape (Quaggin and Kreidberg, 2008). Blood vessel endothelial cells move into the empty region of the lower S and become associated with the layer that will become the podocytes. In that region, an initial capillary loop forms. PDGF receptor mutants result in a glomerulus with a single loop (Alpers et al., 1992). There has been little work on how the capillary branched network forms. However, short cuts could be prevented if there were a barrier positioned perpendicularly in the interior loop (diagram on the far right of Figure 5). For instance, the structure could be precursors to the mesangium. This would allow branching on the sides but not across the barrier, and would naturally lead to the vascular pole restriction.

### A network analysis of flow

Kirchoff’s electric circuit laws are a standard tool for analyzing flows in networks (Newman, 2010). We applied them to the glomerular network where voltage corresponds to pressure, current corresponds to flow, and resistance corresponds to capillary characteristics which impede flow, such as diameter, length, extensibility, etc. We evaluated a simplified model in which all the branches have the same resistance. There was no correction for the filtration that occurs (∼15% of the blood volume is removed during passage through the glomerulus). In this model, the network topology alone determines the local flow rates. The model predicts the direction and magnitude of flow rates in each branch (Figure 7). Since the flow is divided or added at every branch point, it is not surprising that the calculated flow rates are lowest in the branches mid way between the afferent and efferent arterioles. This idealized glomerulus has uniform diameter capillaries, but in the reconstructed glomeruli, the branch diameters vary in relation to the flow rates predicted by connectivity alone (Figure 3C). This is consistent with the idea that capillaries adjust their diameters to adapt to the amount of flow.

### Differences between small and large glomeruli

The minimum number of nodes that must be traversed from afferent to efferent arteriole (the shortest path) was determined for the 12 glomerular networks using standard network theory analysis. Strikingly, this seems relatively independent of glomerular size (Figure 8A). A similar observation was made earlier based on a smaller number of glomeruli from several experimental studies (Wahl et al., 2004). This suggests that glomerulus grows to a certain size, after which, additional branches do not alter the shortest path (i.e., do not make it longer) but instead only make longer alternative paths. When we plotted the shortest paths, they were found to exist almost entirely in the vascular pole side of the glomerulus (Figure 8B). This suggests that paths / later branches are added at the urinary pole.

What would be the stimulus for adding new branches? How would the glomerulus “know” when more branches aren’t needed? Much work indicates that blood vessels adapt to characteristics of fluid flow (Baeyens et al., 2016). Generally, laminar (smooth) flow is thought to be optimized while disrupted / turbulent flow minimized. In the glomerulus, the network topology could determine, to first approximation, the flow rate in the different branches, and the capillaries could adjust their diameter or other properties in order to promote laminar flow. A failure to do this adequately could be the stimulus for formation of new branches.

### Three dimensional histology

The glomerulus is a classic subject of histology, the field of biomedical science that studies the organization of cells in tissues. The histological approach was established by Virchow (1858), and remains an essential part of pathology. Histological considerations are important for regenerative biology which depends crucially on the organization of cells in stem cell niches.

The primary methodology is stained paraffin section, which gives very good views of cell organization. It is surprising though, that histology textbooks illustrate the three dimensional organization of tissues by the use of drawings or diagrams rather than reconstructions. Students are trained to “imagine” the three dimensional structure from single paraffin sections.

This situation is due to the fact that paraffin sections are typically ∼5 *µ*m thick. Because cells are approximately this size, paraffin sections are usually too thick to allow 3D reconstructions with cellular resolution. Additionally, the highest resolution images of paraffin sections are obtained with an oil immersion lens, with an xy resolution of ∼0.5 *µ*m or slightly less. By using the ATUM method to collect 500 nm thick sections, and imaging them with a scanning electron microscope at 50 nm per pixel, the images have 10 times the resolution of a paraffin serial section data in x, y and z dimensions. Thus many of the classic subjects of histology may benefit from three dimensional studies using this approach.

The serial section approach we used is one of several that were developed by neurobiologists to determine synaptic connectivity (Denk and Horstmann, 2004; Micheva and Smith, 2007; Kasthuri et al., 2015;). These methods involved several technical innovations and took advantage of tremendous advances in imaging technology and computer processing power. When used at a lower resolution, the new serial section methods provide an efficient and effective means for investigating biological organization at the cellular level.

## Methods

Three month old C57Bl/6J male mice under anesthesia were fixed by gravity fed cardiac perfusion of Karnovsky fixative (2.5% glutaraldehyde, 1% paraformaldehyde in 100 mM sodium cacodylate, pH 7.4) After several hours in fixative, kidney pieces were treated with 1% osmium, 0.8% potassium ferricyanide in 100 mM sodium cacodylate for 1 hr, in 1% uranyl acetate in water overnight, stained with lead aspartate (Walton, 1979) for 30 min at 60 °C, then dehydrated, infiltrated with epon resin and polymerized. The twelve reconstructed glomeruli were from four mice.

An EM UC-7 ultramicrotome (Leica, Buffalo Grove, IL) with a 6 mm wide histo diamond knife (Diatome, Hatfield, PA) was used cut 500 nm thick sections. Serial sections were collected with an ATUM tape collector (custom built by Ken Hayworth) on kapton tape. The tape was mounted on a silicon wafer (University Wafers, South Boston, MA) then carbon coated (Denton 502B, Moorestown, NJ). 200-250 sections collected.

A Verios (FEI, Hillsboro, OR) or Sigma (Zeiss, Thornwood, NY) field emission scanning electron microscope was used in backscatter mode to collect images. The ATUM approach keeps all the sections, so we searched in the cortex to find glomerulus completely included within series. This is less feasible with block face.

For mapping the sections and image collection with the Verios, the MATLAB program SEM Navigator (kindly provided by Daniel Berger, Harvard University) was used. On the Zeiss, custom Python (http://www.python.org) and Image J (http://imagj.nih.gov) scripts were used to map the sections and Atlas 4 (Fibics Inc., Ottawa, Ontario, Canada) was used to collect the images.

The images were aligned using Linear Stack Alignment with SIFT algorithm of FIJI Image J (https://fiji.sc/). The TrakEM2 module of FIJI was used for segmenting the images. MeshLab (http://meshlab.net) was used for rendering.

It was difficult to document connectivity using the original image data essentially because only one angle of view is available. Keeping track of nodes and connections within the data set was also problematic. We therefore worked with the reconstructions (.obj files) using a software that used virtual reality optics (SyGlass, Morgantown, WV; Pidhorsky et al., 2018). This allowed us to rotate, slice through, and place nodes and connections within the reconstruction. The software exported the node positions and connections for network analysis. The analysis was done in R (https://www.R-project.org) using the igraph (http://igraph.org) and RBGL (http://www.bioconductor.org) packages. The Kirchoff analysis and display (Figure 7) was done in Matlab (http://www.mathworks.com). The display (Figure 8) was made using D3/javascript (http://d3js.org).

The length (number of nodes) of the shortest path was determined by multiplying the adjacency matrix by itself until the afferent and efferent arterioles were connected. Then the adjacency list was used to identify the nodes in the shortest path by tracing backwards from the efferent arteriole.

## Acknowledgements

We thank Michael Saxton for modeling discussions, Boris Slepchenko for help with Kirchoff analysis of flow and Greg Huber for introducing MT to network analysis. We also thank Anthony Chang, Ruchir Trivedi, Ben Afzali, Joe Bonventre, and Robert Colvin for discussions of kidney biology, and Guo Fong for discussions of vascular biology. Special thanks to Patrick La Rivière who made a key observation and to Anthony Chang and Laurinda Jaffe for reading the manuscript. Supported by a grant from the Connecticut Science Fund. J.C.B. was supported by 5T-90DE021989-07 grant from the National Institutes of Health (NIDCR).

